# Winnow-tax: sensitive and precise taxonomic profiling of low-coverage organisms in shotgun metagenomes

**DOI:** 10.64898/2026.10.02.756312

**Authors:** Shan Sun, Anthony A. Fodor

## Abstract

Accurate species-level profiling of shotgun metagenomes becomes difficult when an organism is represented by only sparse sequence coverage. Per-read k-mer classifiers can retain sensitivity under these conditions, but short reads from conserved or shared genomic regions can generate false-positive calls among closely related references. More conservative marker-gene and genome-containment approaches generally provide stronger species-level specificity but may lose sensitivity as genomic sampling becomes sparse. We present winnow-tax, a two-stage taxonomic profiling pipeline that separates sensitive candidate nomination from genome-level confirmation. Candidate species are nominated using sylph, Kraken2, and a branch-rescue procedure, then reads are competitively recruited to a sample-specific reference set. Species presence is evaluated using read support, observed genome breadth, and a Lander-Waterman-based breadth ratio that compares observed breadth with the breadth expected at the measured mean depth. In a controlled synthetic community of 82 genomes with human DNA background, winnow-tax maintained a stronger precision-sensitivity balance than Kraken2/Bracken, sylph, and MetaPhlAn 4 as genome coverage decreased, with its advantage concentrated near the low-coverage detection boundary. In the CAMI III Toy Longitudinal Human Gut benchmark, winnow-tax had higher sensitivity than sylph (0.803 versus 0.672), lower precision (0.923 versus 0.970), and a modestly but significantly higher F1 score (0.858 versus 0.793). In a clinical enteric stool cohort with culture/PCR reference testing, winnow-tax achieved the best performance for *Salmonella*, detecting 33 of 43 culture-positive samples with two false-positive calls (F1 0.846 versus 0.811 for sylph). Together, these results support the advantage of a profiling strategy in which read-level classifications are accepted only when their genomic evidence is distributed as broadly as expected given the sequencing depth.

## Introduction

Shotgun metagenome sequencing has become a widely used approach for characterizing microbial communities because it captures total community DNA present in a sample, rather than targeting a single marker gene. In addition to taxonomic profiles, shotgun metagenomics can provide information on microbial functional genes and allow for genome assembly that cannot be performed with amplicon-based approaches. A central step in analyzing shotgun metagenomes is the classification of sequencing reads against reference databases to determine which microorganisms are present and their relative abundances. Accurate classification is especially important in low microbial biomass samples or shallowly sequenced samples, especially if they are also accompanied by abundant DNA from host or other sources, where the signal-to-noise ratio is low and spurious assignments can obscure biologically meaningful patterns. For example, in human tissue associated datasets, one major challenge is the overwhelming abundance of human DNA relative to microbial DNA. Even after host depletion, residual non-microbial reads may persist because of incomplete reference genomes, human genetic variation, or imperfect filtering workflows (1, 2). The human cancer microbiome provides a well-recognized example of this issue: in tumor tissues, low microbial biomass combined with extensive host background can contribute to false positive microbial detection and has been linked to controversial reports of microbial signatures in cancer and other low-biomass sample types, including placenta and blood (1, 3, 4). Similar challenges arise in environmental samples, where non-microbial background sequences may also be abundant but less well defined. Across these settings, low microbial biomass and low genome coverage place strong demands on both sensitivity and specificity.

In low microbial biomass datasets, only a small fraction of a microbial genome may be sampled, increasing both the difficulty and the uncertainty of taxonomic assignment. Marker gene-based approaches such as MetaPhlAn 4 classify samples using clade-specific marker genes rather than attempting to assign every read (5). By focusing on informative markers, MetaPhlAn 4 can produce taxonomic profiles with high specificity and relatively low false positive rates. However, this strategy requires that reads from diagnostic marker regions are present in the sample. As a result, marker-based methods may have limited sensitivity at very low sequencing depth or low organism abundance, where marker genes may not be sampled even when the organism is present. Similarly, ANI or genome-sketch based approaches such as sylph compare sequencing data to reference genomes using compact k-mer sketches to estimate genome similarity and coverage (6). These methods can provide robust genome-level evidence when sufficient sequence coverage is available, but they may fail to detect taxa when the sampled reads are too sparse to support high similarity or coverage estimates.

In contrast, Kraken2 is a k-mer based per-read classifier that assigns sequencing reads to taxa by matching exact k-mers to a reference database and placing reads using a lowest common ancestor framework (7). Bracken is commonly used downstream of Kraken2 to re-estimate species or genus level abundances from Kraken classifications (8). This approach is computationally efficient and can be highly sensitive because individual reads can contribute taxonomic evidence even when genome coverage is low, but this advantage can come at the cost of increased false positive assignments. One important source of error is the use of compact hash tables and approximate lookup strategies to reduce memory requirements. Kraken2, for example, compresses k-mer hash representations and taxonomic identifiers to enable rapid large-scale classification. Although this design improves computational efficiency, it can also increase the risk of spurious assignments, including those arising from hash collisions or limited taxonomic specificity of short sequence matches. Early Kraken2 releases could classify a read from a single matching minimizer. Later versions require two hit groups under the default settings, which still allows classification from very limited evidence and potentially inflating false positive rates when confidence thresholds are not applied. Kraken2 provides a confidence score cutoff to mitigate this problem, but its use is not always consistent across studies and has not been comprehensively evaluated.

False-positive read assignments are not explained solely by classifier implementation details. Short reads derived from conserved genes, homologous genomic segments, mobile elements, or other sequence shared among related taxa may be compatible with multiple reference genomes. As reference databases expand, dense neighborhoods of closely related genomes can therefore increase ambiguity in species-level assignments (9, 10). A taxon may accumulate many assigned reads while those reads remain concentrated within a restricted part of its reference genome. Increasing the Kraken2 confidence threshold can suppress some weak assignments, but a single classification threshold must trade sensitivity against specificity and does not directly test whether the aggregate evidence is distributed across the candidate genome.

Several previous methods have incorporated more genomic features into taxonomic profiling to reduce false positives. KrakenUniq augments Kraken-style classification with counts of distinct taxon-specific k-mers, providing greater confidence when supporting reads represent many independent sequence features (11). KMCP combines k-mer similarity with approximate genome coverage by dividing reference genomes into chunks and requiring support across those chunks (12). More recently, Wright et al. proposed post-classification confirmation by remapping taxonomically assigned reads to references and evaluating genome coverage (13). These approaches supported the principle that evidence for microorganism presence should not solely depend on the number of reads but also on the distribution across the genome. However, the mechanisms of these methods such as a higher cutoff of genome coverage may not work well in detecting low abundance microbes.

Winnow-tax builds on this principle while targeting the sparse-coverage regime, by using broader candidate nomination to recover low abundance microbes, followed by a presence test to filter out false positives (https://github.com/ssun6/winnow-tax). Candidate taxa are nominated permissively using complementary read- and genome-level signals, after which reads are competitively recruited to a sample-specific set of representative genomes. For each candidate, winnow-tax compares observed genome breadth with the breadth expected at the observed depth under a Lander-Waterman sampling model (14). Thus, a low-abundance organism can pass when a small number of reads are distributed as broadly as expected, whereas a candidate supported by localized cross-mapping can fail despite having a similar read count. We evaluated this framework using a controlled 512-fold genome-coverage series, the CAMI III toy longitudinal human gut benchmark, and a clinical enteric pathogen cohort with culture/PCR reference testing along with Kraken2/Bracken, sylph, MetaPhlAn 4 and KMCP. This approach preserves the sensitivity of methods such as Kraken2 and Bracken while reducing false positives from localized or ambiguous read mapping, providing a practical framework for detecting low-abundance microbial taxa in shotgun metagenomic datasets.

## Results

### Pipeline overview

Winnow-tax was designed to retain the sensitivity of read-level taxonomic classification while reducing false-positive species calls through genome-wide evidence. The pipeline combines candidate taxa identified by sylph and Kraken2 with an additional read-rescue procedure, followed by reference-based evaluation of genome coverage (Figure 1). Kraken2 provides sensitive read-level evidence, but reads with ambiguous k-mer support may be assigned to higher taxonomic ranks when confidence filtering is applied. Winnow-tax re-examines the per-k-mer assignments of these reads and rescues species-level assignments when the k-mer evidence supports an unambiguous descendant of the original classification. These rescued taxa are added to candidates identified by Kraken2 and sylph. Candidates nominated by Kraken2 and branch rescue are then evaluated using genome-wide alignment evidence. Rather than accepting a species based on read count alone, winnow-tax compares the observed breadth of genome coverage with that expected from its estimated coverage. Taxa with insufficient or highly localized support are removed. The resulting workflow therefore uses read-level evidence to maximize candidate recovery while requiring genome-level consistency before accepting species detected by read-based classification.

**Figure 1.**
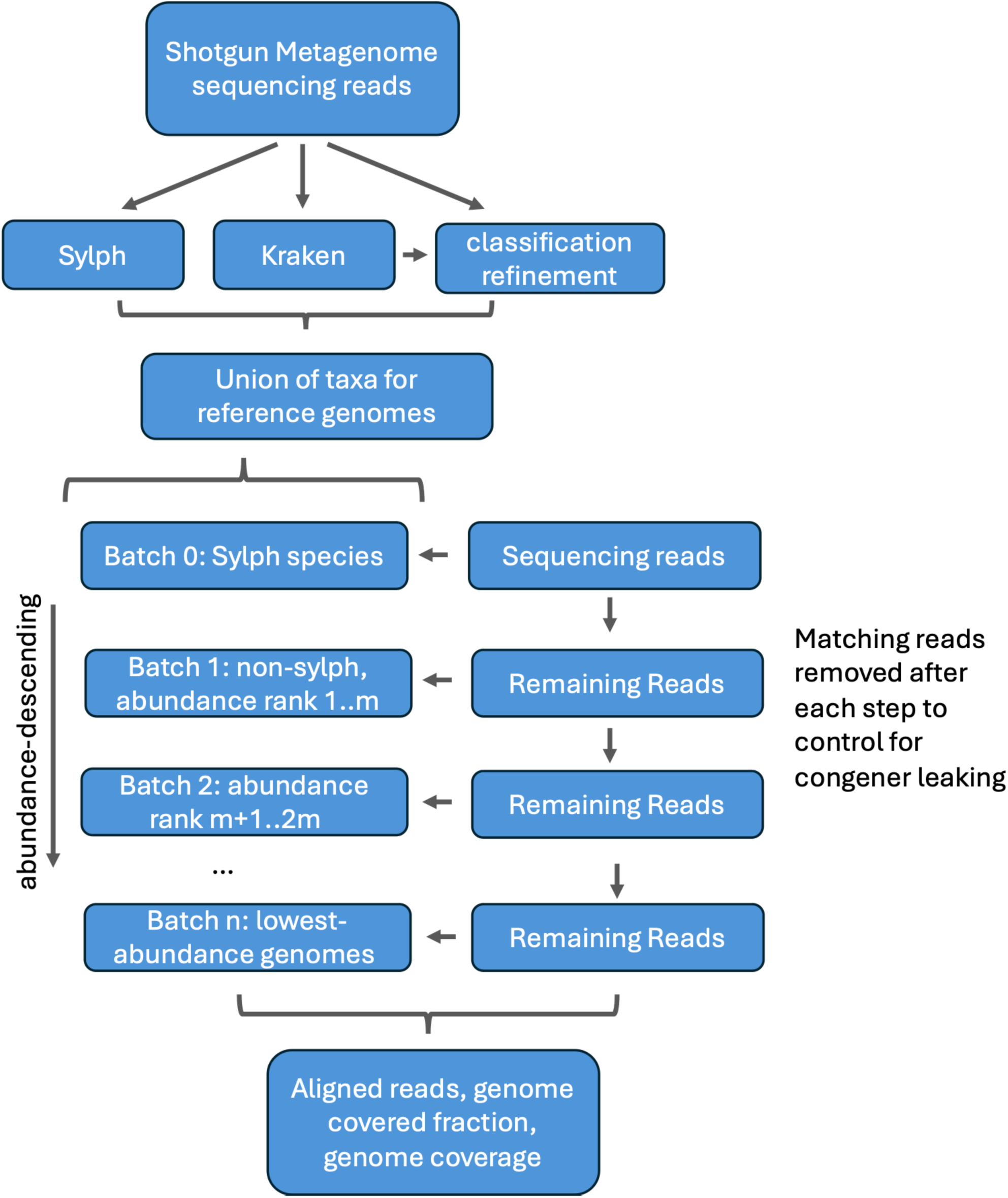
Overview of the winnow-tax pipeline. Candidate species are identified using sylph containment profiling, Kraken2 classification, and read-level taxonomic rescue. Candidate genomes nominated by Kraken2 and read-level rescue are subsequently evaluated by competitive read alignment and genome-wide coverage consistency to retain well-supported species while filtering likely false-positive detections.

### Winnow-tax retains species detection as genome coverage falls

To benchmark the performance of winnow-tax, sylph, Kraken2, MetaPhlAn 4 and KMCP across a range of samples with varied microbial coverages, we constructed a mock community of 82 known genomes (15) and simulated shotgun metagenomic reads across a 10-step 2-fold decreasing gradient of genome coverage levels from 2.5× to 0.00488×, with human genomic reads added as a background noise. We used both GTDB and RefSeq databases for benchmarking to analyze if their performance varied between databases. Because MetaPhlAn 4 uses a fixed marker-gene database that reports NCBI taxonomy and cannot be customized, it was included only in the RefSeq/NCBI benchmark.

Across the 512-fold change coverage mock community profiled using GTDB database, winnow-tax achieved the highest mean F1 score of all methods and parameter settings evaluated (F1 = 0.949), with mean precision of 0.931 and recall of 0.975. The competing methods generally exhibited a trade-off between sensitivity and specificity. Sylph achieved near-perfect precision (0.998) but substantially lower recall (0.694), resulting in a mean F1 of 0.719. In contrast, Bracken maintained near-perfect recall across confidence thresholds (0.989-1.000) but showed markedly lower precision (0.083-0.341 for confidence 0.1-0.9), with a maximum F1 of 0.506 at the best performing confidence threshold tested. KMCP spanned this trade-off depending on setting, with the best of the three settings compared (m1) reaching a mean F1 of 0.774 (precision = 0.640, recall = 0.998), but no KMCP setting matched the combined precision and recall achieved by winnow-tax.

The principal advantage of winnow-tax over sylph was improved sensitivity at low sequencing coverage. Sylph recall declined sharply below approximately 0.0391× coverage, reaching 0.160 at 0.0195×, 0.025 at 0.00977×, and 0 at 0.00488×. In comparison, winnow-tax maintained recall above 0.85 across the full dilution series. Conversely, the principal advantage of winnow-tax over Bracken was substantially higher precision. Even at Bracken’s most stringent confidence threshold 0.9, precision was 0.341 compared with 0.931 for winnow-tax. Because Bracken recall remained near 1.0 across all tested thresholds, its lower F1 scores were driven primarily by false-positive classifications. KMCP performance varied substantially among parameter settings. The more permissive setting m1 achieved the highest F1 by maximizing recall at the expense of precision, whereas the more stringent settings m3 and m5 increased precision but reduced sensitivity for low coverage samples.

Abundance-estimation accuracy showed a similar pattern. Winnow-tax had the lowest mean L1 distance (0.125, compared with 0.476 for sylph, 0.411-0.985 across KMCP settings, and 0.85-1.35 across Bracken confidence thresholds 0.1-0.9), with its advantage concentrated at low coverage: below approximately 0.04×, sylph and KMCP errors rose sharply, whereas winnow-tax L1 remained low. At higher coverage, sylph achieved similar or slightly lower L1 than winnow-tax.

### Winnow-tax improved species detection in complex communities

Winnow-tax was also tested on the new CAMI III toy human gut benchmark dataset (16), consisting of 20 samples. To keep the nomenclature consistent, we classified the 1,619 bacterial source genomes with GTDB-Tk (17) and evaluated results in GTDB R226 taxonomy. This approach avoided the NCBI-to-GTDB nomenclature mismatch that can confound direct comparisons. Across the 20 CAMI III samples evaluated against GTDB, winnow-tax achieved the highest mean F1 score (0.858) among all methods and parameter settings tested. Paired comparisons across samples confirmed that the performance advantage of winnow-tax was statistically significant. In contrast to the varied coverage mock communities, the CAMI III benchmark represents more complex and realistic microbial communities, providing a test of performance across a broader range of taxa, abundances, and community compositions.

The closest-performing method was sylph, with a mean F1 of 0.793 compared with 0.858 for winnow-tax (paired t-test, P<1e-10). The improvement of winnow-tax over sylph was driven primarily by higher recall (0.803 versus 0.672), with a modest reduction in precision (0.923 versus 0.970). Winnow-tax also substantially outperformed all KMCP and Bracken settings in F1 score. Compared with the best-performing KMCP setting (m5), winnow-tax increased precision from 0.523 to 0.923 and recall from 0.603 to 0.803, resulting in an F1 increase from 0.559 to 0.858 (paired t-test, P<1e-15). Similarly, Winnow-tax outperformed Bracken with its best performing confidence cutoff, with a mean F1 of 0.858 compared to 0.374 from Bracken (paired t-test, P<1e-20). Relative to Bracken, winnow-tax maintained substantially higher precision while retaining high recall, with precision of 0.923 compared to 0.237, and recall decreased only modestly from 0.884 to 0.803.

Winnow-tax also provided accurate abundance estimates. Its mean L1 error was 0.101, identical to sylph and substantially lower than all KMCP (0.346-0.446) and Bracken (0.435-0.772 for confidence 0.1-0.9) settings. Winnow-tax therefore combined the abundance accuracy of sylph with substantially greater species-level sensitivity, while providing considerably more accurate community profiles than KMCP or Bracken. The per-species predicted-versus-true abundance plots (Figure 3E) further illustrate these differences. Winnow-tax produced fewer false-negative species-sample pairs than sylph (19.7% versus 32.9%), while maintaining a relatively low false-positive fraction (7.6% versus 3.0%). Importantly, many of the additional species recovered by winnow-tax occurred at low true abundance and were nevertheless assigned predicted abundances close to the identity line (Figure 3E, winnow-tax panel, highlight in red rectangle). Thus, the increased sensitivity of winnow-tax was not limited to detecting additional low-abundance taxa, but extended to accurate quantification of many taxa that were missed by sylph. In contrast, Bracken/Kraken2 and KMCP profiles were more dispersed around the identity line with larger deviations across different parameter settings, indicating less accurate abundance estimation (Figure 3E). Together, these results show that the sensitivity and precision observed in the simpler varied coverage series extend to more complex and realistic microbial communities, with winnow-tax providing improved recovery and quantification of low-abundance species without the large increase in false-positive detections observed for the other methods.

**Figure 2.**
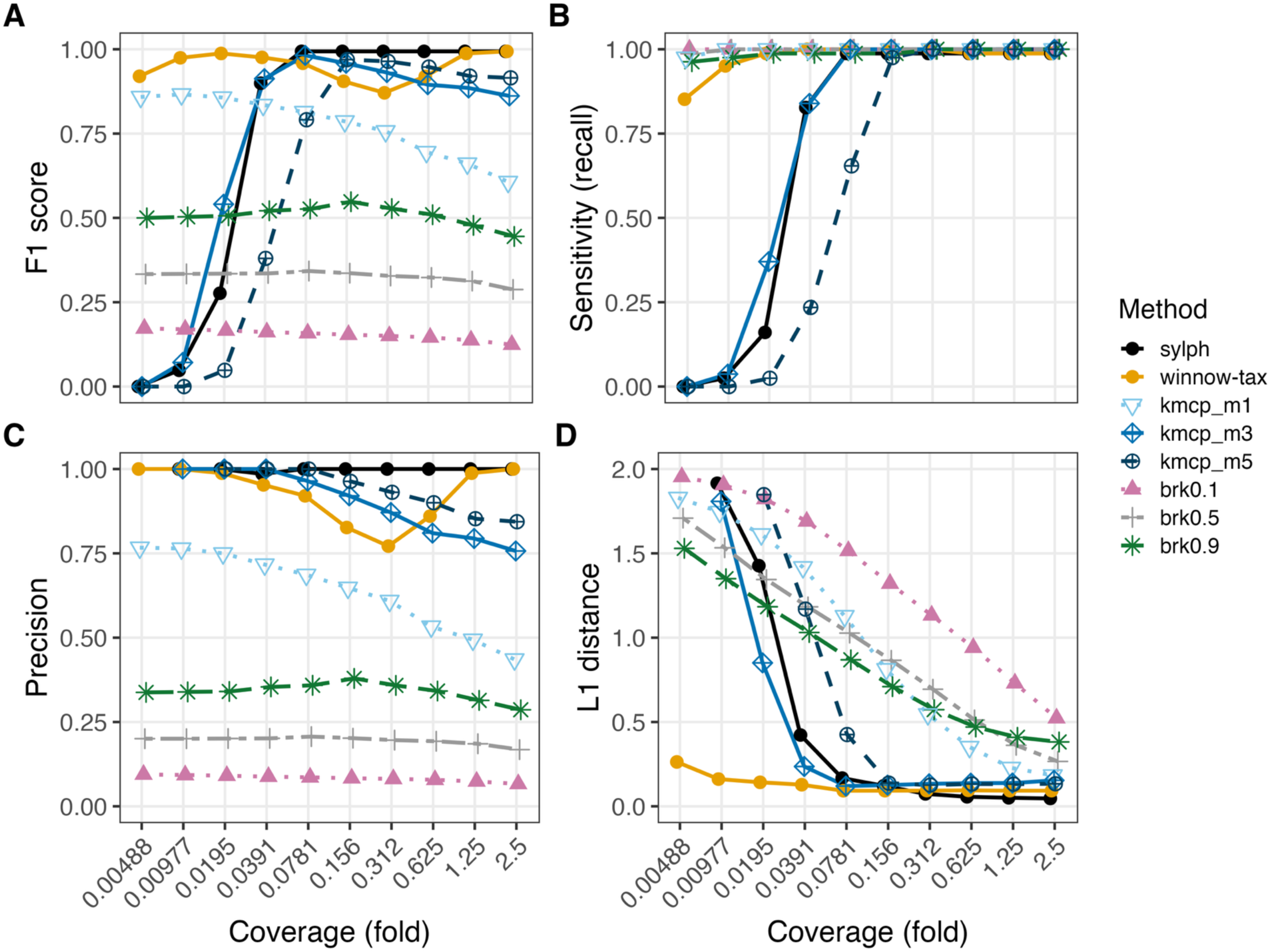
Winnow-tax maintains species detection across a 512-fold coverage range. Ten simulated samples containing 82 bacterial and archaeal genomes and human genome were profiled across per-genome coverages from 2.5× to 0.00488× with winnow-tax, Kraken2/Bracken, sylph and KMCP using GTDB database. (A) F1 score, (B) sensitivity, (C) precision, and (D) L1 distance between predicted and true relative abundance. Winnow-tax retained high recall at low coverage while maintaining substantially higher precision than Bracken. Sylph remained highly precise but lost sensitivity at the lowest coverages. In panel D, samples in which a method reported no species are omitted, and this removes one sample each for sylph and KMCP mode 3 and two samples for KMCP mode 5. brkX indicates Bracken applied to Kraken2 classifications at confidence X; kmcp_mX indicates KMCP profiling mode X.

**Figure 3.**
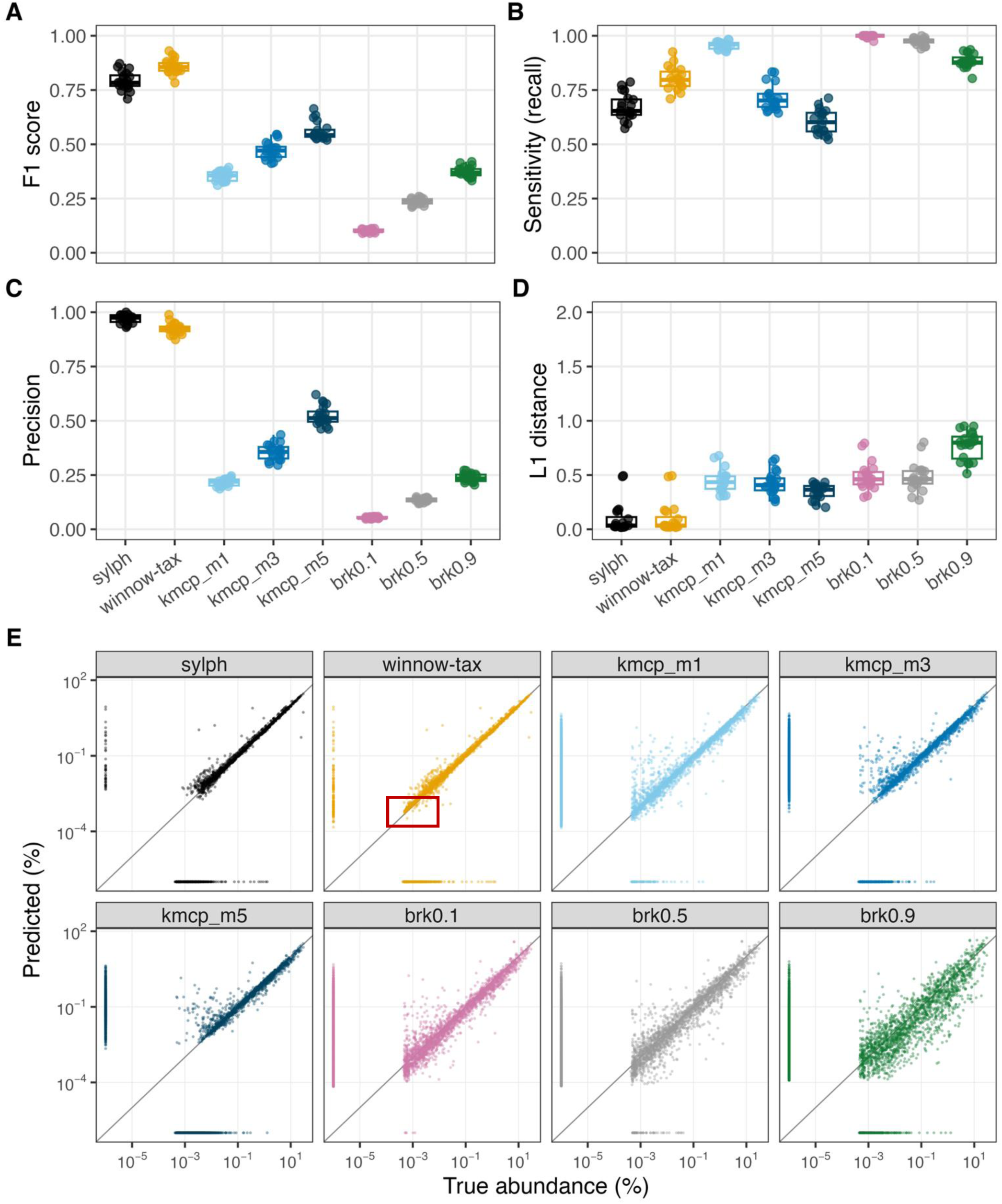
Winnow-tax improves species recovery in complex human-gut communities. Twenty CAMI III human-gut samples were profiled with winnow-tax, Kraken2/Bracken, sylph and KMCP using GTDB database. (A) F1 score, (B) sensitivity, (C) precision, (D) L1 distance, and (E) predicted versus true relative abundance across species. Boxes show medians and interquartile ranges with individual samples overlaid. Winnow-tax achieved the highest F1 by combining higher recall than sylph with substantially higher precision than Bracken and KMCP. In panel E, marginal stripes represent false-positive and false-negative species-sample pairs. Boxed area highlights low-abundance taxa detected by winnow-tax but missed by sylph. brkX indicates Bracken applied to Kraken2 classifications at confidence X; kmcp_mX indicates KMCP profiling mode X.

### Winnow-tax improves pathogen detection in clinical stool metagenomes

We evaluated pathogen detection on the Canadian enteric stool metagenome cohort from Peterson et al. (18), which includes shotgun metagenomic sequencing of clinical stool specimens with paired routine diagnostic testing for bacterial enteric pathogens. Diagnostic labels were derived from culture and PCR-based reference testing. When Salmonella and Campylobacter were considered detected using culture-confirmed infection status as the reference standard, winnow-tax detected more culture-positive samples than sylph for both target organisms while maintaining similar specificity. For Salmonella, winnow-tax detected 33 true positives compared with 30 for sylph, resulting in a higher F1 score (0.846 versus 0.811). For Campylobacter, winnow-tax detected 48 true positives compared with 46 for sylph, with F1 scores of 0.835 and 0.821, respectively.

The primary advantage of winnow-tax over Kraken2/Bracken was substantially higher precision, most clearly for Salmonella. For Salmonella, winnow-tax achieved a precision of 0.943, compared with 0.237 for Bracken at Kraken2 confidence 0.1 and 0.398 at confidence 0.5. These differences corresponded to only 2 false-positive detections across 322 sequencing runs for winnow-tax, compared with 135 and 59, respectively, for the two Bracken settings. Increasing the Kraken2 confidence threshold reduced false-positive detections but also reduced sensitivity. At confidence 0.9 Bracken precision reached only 0.494 and still produced 39 false-positive detections. Campylobacter behaved differently, where Bracken retained higher sensitivity (0.932 and 0.949 at confidences 0.1 and 0.5) than winnow-tax (0.814), at a modest cost in precision (0.764-0.767 versus 0.857), giving it a slightly higher F1 (0.840-0.848 versus 0.835). The precision advantage of winnow-tax over Bracken was therefore concentrated in Salmonella, where false-positive calls were an order of magnitude more frequent.

The same overall pattern was observed when PCR-based and composite reference standards were used instead of culture-confirmed status, indicating that the relative performance of the methods was not dependent on the clinical reference standard. KMCP was not evaluated on this cohort because of the extensive time and resources required to run KMCP on over 300 samples. Shigella was also tested in that dataset, but they are pathotype-defined within the Escherichia coli/Shigella complex and cannot be resolved reliably by species-level taxonomic profiling alone.

### Winnow-tax performance is maintained with a RefSeq reference database

To determine whether the observed performance depended on the underlying reference taxonomy, we repeated the benchmarks using RefSeq. We first compared winnow-tax performance under GTDB and RefSeq on both mock-community benchmarks. GTDB produced higher F1 scores on both mock-community benchmarks: 0.949 versus 0.881 on the varied coverage series and 0.858 versus 0.749 on CAMI III (Fig. 5A and B). For abundance estimation, the difference was larger with L1 distance decreasing from 0.306 under RefSeq to 0.125 under GTDB on the spike-in series and from 0.528 to 0.101 on CAMI III (Fig. 5C and D). In contrast, reference database choice had little effect on the clinical cohort. For Salmonella, F1 was 0.846 with GTDB and 0.842 with RefSeq, while Campylobacter F1 was identical at 0.835 (Fig. 5E). Across all 322 sequencing runs of clinical samples, the RefSeq analysis differed from the GTDB analysis by only one true-positive and one false-positive call.

**Figure 4.**
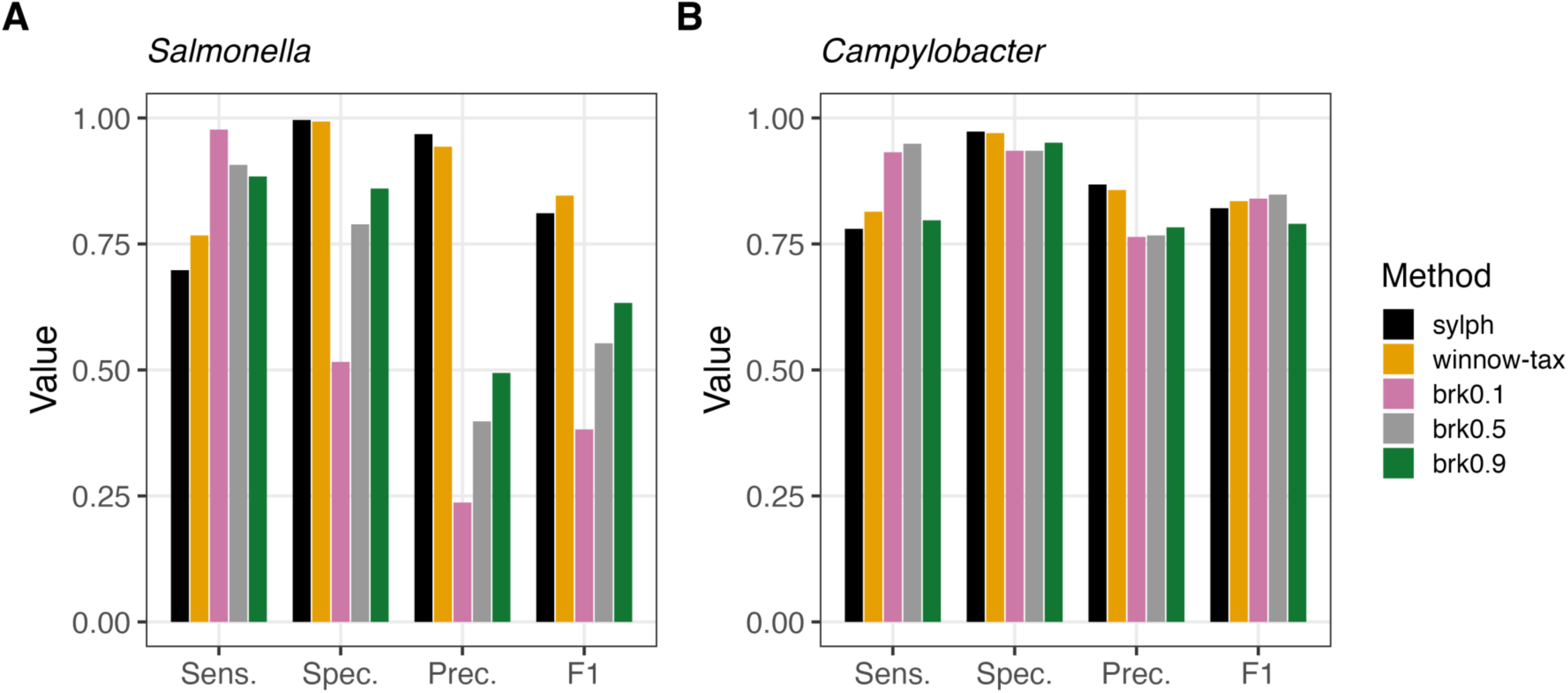
Detection of enteric pathogens in clinical stool metagenomes. Performance for detection of *Salmonella* (A) and *Campylobacter* (B) was evaluated against culture-confirmed status in 322 sequencing runs of clinical stool metagenomes using GTDB-based profiles. Sensitivity, specificity, precision, and F1 are shown for all methods. Winnow-tax detected more true-positive samples than sylph for both pathogens while maintaining similar specificity. Bracken is shown at Kraken2 confidences 0.1, 0.5 and 0.9. Relative to Bracken, winnow-tax substantially reduced false-positive *Salmonella* calls while retaining high sensitivity, whereas in panel B the two were comparable and Bracken achieved a slightly higher F1 at the two lower confidences. brkX indicates Bracken applied to Kraken2 classifications at confidence X.

**Figure 5.**
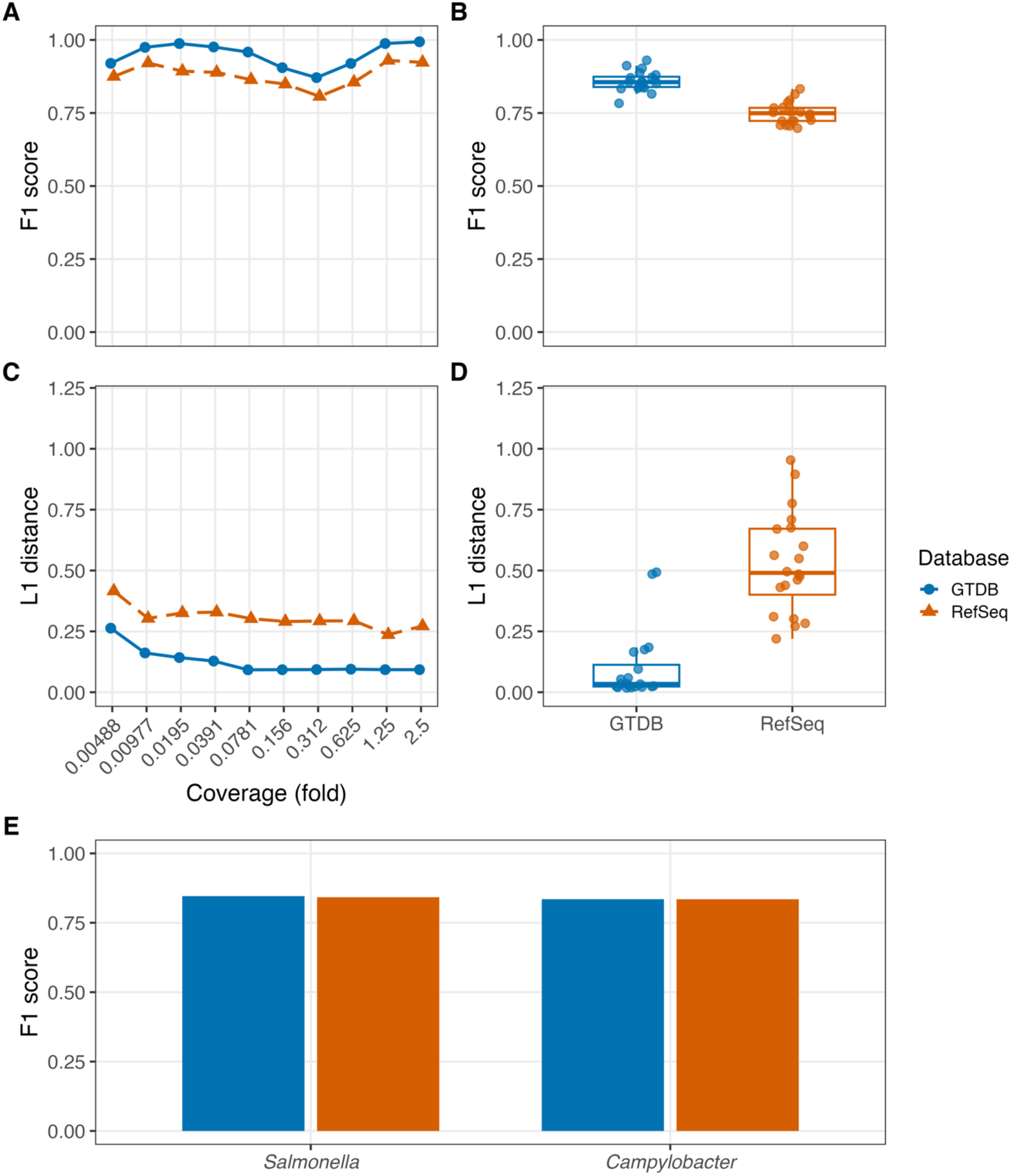
Effect of reference database on winnow-tax performance. Winnow-tax was evaluated using GTDB and RefSeq across the 512-fold coverage, CAMI III, and clinical datasets. (A) 512-fold coverage F1, (B) CAMI III F1, (C) 512-fold coverage L1 distance, (D) CAMI III L1 distance, and (E) clinical pathogen F1. GTDB produced higher detection and abundance accuracy on both mock communities, whereas clinical *Salmonella* and *Campylobacter* performance was nearly unchanged.

Because MetaPhlAn 4 reports NCBI taxonomy and its marker database cannot be customized, we included its performance comparison in this RefSeq benchmark with other databases. Winnow-tax achieved the highest F1 score on both mock-community datasets: varied genome coverage benchmark and CAMI III. The overall patterns observed for sylph and Kraken2/Bracken were consistent with those seen under GTDB (Figures 2 and 3). Sylph maintained high precision but was limited by recall, whereas Bracken retained high recall across confidence thresholds (0.910-0.930 on the coverage series and 0.777-0.866 on CAMI III) at the cost of substantially lower precision.

MetaPhlAn 4 showed a precision-recall profile similar to that of sylph. At the three lowest genome coverage levels, recall was 0.000, 0.000, and 0.063 for MetaPhlAn 4 and 0.000, 0.013, and 0.125 for sylph. In contrast, winnow-tax maintained recalls of 0.787, 0.875, and 0.887 across the same samples. Bracken also retained high recall (0.887, 0.912, and 0.912), but with substantially lower precision. Thus, although MetaPhlAn 4 and sylph use fundamentally different profiling strategies, both became recall-limited at very low coverage, whereas winnow-tax retained much of the low-abundance signal without the large increase in false-positive detections observed with Bracken. On CAMI III, sylph and MetaPhlAn 4 had higher precision than winnow-tax (0.858 and 0.882 versus 0.845) but lower recall (0.557 and 0.504, respectively, versus 0.674). Winnow-tax therefore achieved higher F1 primarily by recovering a larger fraction of the true community at a small cost in precision. By comparison, no Bracken setting exceeded a precision of 0.483 on either mock-community dataset despite its consistently high recall.

On the clinical cohort, a similar precision-recall pattern was observed. For Salmonella, winnow-tax, sylph, and MetaPhlAn 4 each produced only one false-positive detection across 322 sequencing runs, resulting in nearly identical precision (0.970, 0.968, and 0.967, respectively). Their performance differences were therefore driven by sensitivity: winnow-tax detected 32 true-positive samples, compared with 30 for sylph and 29 for MetaPhlAn 4, yielding F1 scores of 0.842, 0.811, and 0.795, respectively. In contrast, all Bracken settings had substantially lower precision, with a maximum of 0.582 at confidence 0.9 among the settings shown. Campylobacter was the only clinical comparison in which a Bracken setting achieved the highest F1 score. MetaPhlAn 4 had the highest precision (0.870; 6 false positives) but detected only 40 of 59 culture-positive samples, resulting in an F1 score of 0.762, compared with 0.835 for winnow-tax and 0.821 for sylph. Bracken at confidence 0.5 achieved the highest F1 (0.855), as its high recall (0.949) outweighed its lower precision (0.778), but overall the performances of sylph, winnow-tax and Bracken are similar for Campylobacter, with F1 ranging from 0.821 to 0.855.

### Computational cost

Runtime was evaluated using CAMI III samples containing a mean of 33.27 million reads. Ranked by CPU time per million reads (GTDB databases for sylph, Kraken2, winnow-tax and KMCP; MetaPhlAn 4 with its own marker database), sylph was the least computationally intensive at 3.2 CPU-s per million reads, followed by Kraken2 at 33.5 CPU-s, Winnow-tax at 108.2 CPU-s and MetaPhlAn 4 at 218.8 CPU-s, whereas KMCP had by far the highest computational requirement at 31,429 CPU-s per million reads, 2 to 4 orders of magnitude higher than other methods (Table 1). At the CAMI III sequencing depth, this corresponds to approximately 1,045,779 CPU-s, or 290.5 CPU-h (12.1 CPU-days), for a single 33-million-read sample. KMCP therefore required approximately 291-fold more CPU time than winnow-tax while not exceeding its profiling accuracy on either mock-community benchmark, representing a substantial computational disadvantage for large-scale applications. Switching from GTDB to RefSeq reduced CPU use by 17% for winnow-tax, 29% for sylph and 40% for Kraken2. The substantially smaller RefSeq reference also reduces storage and lowers memory requirements, making it a practical alternative for samples dominated by well-characterized taxa and without a large representation of novel MAGs.

**Table 1.** Computational resources and runtime of taxonomic profiling methods. Wall-clock time, CPU time, and database size were measured for 20 CAMI III samples containing 33.27 million individual reads on average. Runtime is reported per sample and normalized per million reads. CPU time was used for cross-method comparisons because thread allocations differed among tools. Database sizes are reported in GiB.

| Tool | Database | DB size | Cores | Wall time/sample (s) | CPU time/sample (s) | Wall time/1M reads (s) | CPU time/1M reads (s) |
| --- | --- | --- | --- | --- | --- | --- | --- |
| sylph | RefSeq | 3.8 GiB | 16 | 68.0 | 76.2 | 2.0 | 2.3 |
| sylph | GTDB | 17 GiB | 16 | 74.6 | 106.8 | 2.2 | 3.2 |
| Kraken2 | RefSeq | 147 GiB | 16 | 209.0 | 666.6 | 6.3 | 20.0 |
| Kraken2 | GTDB | 665 GiB | 16 | 1,056.0 | 1,115.7 | 31.7 | 33.5 |
| winnow-tax | RefSeq | 179 GiB | 16 | 1,381.1 | 2,974.3 | 41.5 | 89.4 |
| winnow-tax | GTDB | 809 GiB | 16 | 2,327.2 | 3,599.3 | 69.9 | 108.2 |
| MetaPhlAn 4 | vJan25-SGB | 33 GiB | 12 | 774.8 | 7,280.3 | 23.3 | 218.8 |
| KMCP | GTDB | 159 GiB | 32 | 37,708.1 | 1,045,779.1 | 1,133.3 | 31,429.2 |

## Discussion

Species-level taxonomic profiling of shotgun metagenomes requires balancing two competing goals: detecting organisms represented by relatively few reads while avoiding false-positive assignments caused by conserved sequence, closely related genomes, and incomplete representation of microbial diversity in the reference database. This trade-off has been evident across independent metagenomic benchmarking efforts, including the CAMI challenges, in which performance generally deteriorates toward lower taxonomic ranks and varies substantially with both method and parameter choice (19, 20). Previous benchmarking studies have similarly shown that no single profiling strategy consistently maximizes both sensitivity and specificity across datasets (21, 22).

The results presented here suggest that winnow-tax shifts this sensitivity-specificity trade-off by combining complementary forms of evidence rather than relying exclusively on either read-level classification or genome-level containment. Kraken2 achieves high sensitivity by assigning individual reads using minimizer-based taxonomic evidence, and Bracken subsequently redistributes those assignments to estimate species abundance. This strategy preserves weak signals but can assign reads from conserved genomic regions to closely related taxa. At the other extreme, sylph uses genome-to-metagenome containment and ANI-based filtering, providing high species-level precision with very low computational requirements. MetaPhlAn 4 similarly emphasizes specificity by requiring support from species-level marker genes rather than classifying the complete read set.

Winnow-tax appears to benefit from using these signals sequentially: permissive read-level evidence identifies candidate organisms, whereas the subsequent genome-level coverage assessment tests whether the supporting reads are distributed in a way consistent with the presence of the candidate genome. This distinction is important because the method is not simply averaging the outputs of two profilers. Rather, one source of evidence supplies sensitivity and a second, partially independent source provides a biological consistency check. The resulting profile retained much of the low-coverage sensitivity of Kraken2/Bracken while remaining much closer to sylph in false-positive abundance.

A related strategy has independently emerged in KMCP, which supplements k-mer matching with evidence of coverage across genome chunks specifically to reduce false-positive taxonomic assignments (12). The fact that two independently developed methods use genome-distributed evidence to regularize k-mer-based detection supports a broader principle: at species level, the spatial distribution of sequence support across a genome can provide information that is not captured by the number of matching reads alone. In the present benchmarks, winnow-tax achieved better balance with substantially lower computational requirements than KMCP.

The largest advantage of this strategy was observed at low genome coverage. In the dilution series, differences between winnow-tax and sylph were concentrated below approximately 0.04-fold coverage, and the CAMI community results similarly showed that many of the additional species recovered by winnow-tax occurred at the low-abundance end of the distribution. This regime is intrinsically difficult because the number of observable taxon-specific k-mers or marker genes becomes increasingly sparse as sequencing depth decreases. Sylph explicitly addresses this problem by modeling k-mer containment under low coverage using zero-inflated Poisson statistics, whereas MetaPhlAn 4 relies on coverage of clade-specific marker sequences (5, 6). Our results suggest that retaining read-level candidate evidence before requiring genome-level consistency can extend useful detection further into this sparse-sampling regime.

Importantly, the additional sensitivity was not simply obtained by reporting more low-abundance taxa. In the CAMI abundance plots, many taxa detected by winnow-tax but missed by sylph remained close to the predicted-versus-true abundance identity line. Thus, the principal gain was recovery of additional quantitatively supported taxa rather than indiscriminate expansion of the reported community. This distinction is important because low-abundance false positives are a recognized weakness of metagenomic classifiers and can be poorly reflected by aggregate abundance-distance metrics alone (21).

The clinical stool cohort provides a complementary test of this behavior because the objective is no longer reconstruction of an entire community but detection of a specific organism against an independent clinical reference standard. For *Salmonella*, winnow-tax, sylph, and MetaPhlAn 4 all maintained very high precision, but winnow-tax recovered additional culture-positive samples. In contrast, less stringent Kraken2/Bracken settings recovered many true positives but also generated substantially more false-positive detections. This result extends the mock-community findings to a setting in which false-positive calls have a direct interpretive consequence. False-positive control is particularly important in clinical metagenomics. Unlike targeted microbiological assays, shotgun sequencing simultaneously interrogates a very large sequence space, and spurious alignments, contaminated reference sequences, closely related organisms, and environmental or reagent sequences can all generate apparently plausible pathogen signals. Consequently, clinical metagenomic workflows require careful database curation, orthogonal validation, and thresholds that distinguish low-level background from clinically meaningful detection (23, 24). CAMI II similarly found pathogen identification to be an area in which reproducibility among metagenomic methods remained challenging (19).

The *Campylobacter* results also provide an important counterpoint to the overall benchmark. Bracken at confidence 0.5 achieved the highest F1 for this target because its higher recall compensated for lower precision. However, the higher *Campylobacter* recall of Kraken2/Bracken should also be interpreted in the context of the sequence evidence supporting these detections. Several culture-positive samples detected by Kraken2/Bracken but not by winnow-tax contained insufficient genome-wide evidence to satisfy the winnow-tax coverage criterion. Culture positivity indicates that *Campylobacter* was present in these samples, but it does not necessarily imply that the sequencing depth was sufficient to generate robust metagenomic evidence of the organism. A permissive read-level classifier can make a positive call from a small number of matching reads, and in a culture-positive sample such a call will be scored as a true positive even when the supporting reads are sparse or restricted to a small portion of the genome. Thus, some of the additional recall achieved by Kraken2/Bracken may reflect stochastic recovery of a small number of informative reads rather than consistently recoverable genome-level evidence.

Reference-based metagenomic profiling depends strongly on database composition and taxonomy. Database choice can alter both sensitivity and taxonomic accuracy, particularly for poorly represented or uncultivated organisms (22, 25, 26). More complete databases can improve sensitivity but may also increase ambiguity among closely related genomes and computational cost. Our GTDB-RefSeq comparison further shows that database content and taxonomic structure are difficult to separate. GTDB uses a genome-based phylogeny with ANI-defined species clusters, including placeholder species for unnamed genomes, whereas NCBI Taxonomy incorporates formal nomenclature, synonyms, and incompletely resolved taxa (27, 28). Consequently, GTDB may be advantageous for broad community profiling and poorly characterized diversity, whereas the smaller RefSeq reference can be practical for well-characterized taxa with lower storage and memory requirements. Environment-specific isolates or MAGs may further improve performance when standard references are incomplete (26).

In addition to database size, the approximately four-order-of-magnitude range in CPU requirements observed also emphasized that computational resources are also an important consideration when selecting a profiling strategy. Winnow-tax occupies an intermediate position: it is more computationally demanding than sylph or Kraken2 but less than MetaPhlAn 4 in this benchmark and far less intensive than KMCP. This trade-off may matter differently across applications. For large epidemiological cohorts in which the organisms of interest are reasonably abundant, sylph’s computational efficiency may outweigh the additional low-coverage sensitivity provided by winnow-tax. Conversely, low-biomass studies or analyses in which rare organisms are scientifically important may justify additional computation required by winnow-tax to improve recall.

A key strength of winnow-tax is the interpretability of its filtering framework. Candidate taxa are evaluated using explicit criteria based on read support, genome coverage breadth, and the coverage-normalized breadth ratio rather than a single score. This makes failures interpretable, because taxa may be rejected due to insufficient reads, fragmented genome coverage, or poor reference representation. These same features impose limitations. Winnow-tax remains reference dependent and may miss organisms absent or divergent from available genomes present in the reference database. Its requirement for genomic support can also reduce sensitivity near the metagenomic detection limit, where a true organism may be represented by only a few reads. In addition, species-level classification cannot resolve pathotypes defined by mobile or accessory genes, such as Shiga toxin-producing *Escherichia coli*, without complementary gene-level analysis.

## Conclusions

Winnow-tax addresses a recurring problem in species-level metagenomic profiling: highly sensitive read-level classifiers retain weak signals but can generate false-positive taxa, whereas more restrictive genome- or marker-based approaches provide strong specificity but may lose organisms when sequence coverage becomes sparse. By using read-level evidence to nominate candidates and genome-level coverage evidence to evaluate those candidates, winnow-tax recovered substantially more low-abundance taxa than the containment-based and marker-based profilers evaluated here while retaining much of their false-positive control. The same balance was observed in a complex simulated community and, for pathogen detection in clinical stool metagenomes. Overall, winnow-tax provides taxonomic classification with high sensitivity balanced with better controls for false positives, and is particularly useful for recovery of low-abundance species from shotgun metagenomes.

## Methods

### Study design

Shotgun metagenome reads were adapter- and quality-trimmed using fastp (29). Reads were classified using three complementary strategies to recover candidate taxa (Figure 1). First, sylph is run against the GTDB R226 database, retaining all containment hits without an abundance threshold. Second, Kraken2 is run against a GTDB R226 representative-genome database with a confidence score cutoff of 0.7. Third, a branch-rescue procedure re-examines the confidence-independent per-read k-mer profile and reassigns reads that Kraken2 collapsed to higher taxonomic ranks back down to species. In general, Kraken2 assigns reads to the lowest common ancestor (LCA) of their k-mer hits, ambiguous reads may therefore lose species-level assignments at higher confidence thresholds. To recover this signal without lowering the confidence threshold, we re-examined each read’s per-k-mer taxonomic assignments and reassigned reads only to more specific descendants within their original lineage. Two rescue rules were applied sequentially. First, under a monophyletic-descent rule, a read was reassigned to its deepest supported taxon when all other k-mer hits were ancestral to that taxon, the original Kraken2 call was also ancestral, and the deepest taxon was supported by at least 8 k-mers. Reads with weaker or conflicting lineage support were unchanged. Second, for unreassigned genus-level reads, a dominant-species rule counted species-level k-mer hits within the assigned genus. A read was reassigned to the uniquely most-supported species when at least two congeneric species were represented and the dominant species accounted for at least 10% of all k-mers (confidence score `≥` 0.1) used in the Kraken2 confidence denominator. Ties and sub-threshold reads remained at the genus level. Rescued species were added to the sample-specific candidate reference set. The union of nominated species by these three steps defines a sample-specific reference set of representative genomes.

Reads are then recruited to the sample-specific reference using minimap2 (30) in a cascade style to minimize the chance of reads assigned to congeners. Genomes detected by sylph claim their reads first in an order of descending abundance. Claimed reads were then removed from the pool for remaining steps. The genomes detected by kraken and branch-rescue then compete for the leftover reads in abundance-ordered batches (default 10 batches), with each read assigned to a single genome. Alignments are filtered using minimum thresholds of 95% read identity and 80% read coverage. For each genome, winnow-tax calculates the observed covered fraction, mean depth, read count, and the expected covered fraction under the Lander-Waterman model, B_expected = 1 − exp(−D), where D is mean depth across the full reference genome. The breadth ratio was defined as B_observed/B_expected. By default, a genome is called present if the ratio of observed to expected covered fraction, referred to as the breadth ratio, is at least a specified threshold R (default 0.85), the covered fraction is at least C (default 0.001), and at least 50 reads are recruited. Genomes that fail these criteria are classified as suspicious, because the reads likely accumulated over a limited genomic region, or with insufficient read support for covered fraction of the genome. Because sylph calls are already supported by k-mer containment distributed across the genome, they are admitted to the final profile on that evidence and are not subject to the read-count, breadth-ratio and covered-fraction gates. The gates therefore apply to candidates nominated by Kraken2 and branch-rescue. Thus, winnow-tax uses the breadth ratio, rather than a fixed covered-fraction cutoff alone, as its core presence criterion for candidates nominated by Kraken2 and branch-rescue. This design allows genuine low-abundance organisms to be detected when reads are distributed as expected across the genome, while rejecting cross-mapping artifacts that recruit reads to only a small portion of a genome.

### Benchmark datasets

We evaluated species-level taxonomic profiling across three datasets: a controlled varied genomic coverage series with a range from 2.5× to 0.00488×, with a 10-step 2-fold decreasing gradient, a complex simulated human gut community, and a clinical stool metagenomic cohort. The benchmark compared winnow-tax with sylph, Kraken2/Bracken, KMCP, and MetaPhlAn 4 and evaluated species detection, abundance estimation, computational cost, and robustness to reference database choice.

For the decreasing coverage benchmark, a synthetic community of 82 microbial genomes (61 bacterial and 21 archaeal) was constructed based on the MOCK2 genome set described by Meslier et al. (15). Genomes in the list but no longer in RefSeq database were not included. Ten samples were simulated using art_illumina (31) with the same organisms at progressively decreasing abundance, with per-genome coverage halved between consecutive samples from approximately 2.5× to 0.00488×, spanning a 512-fold range. Microbial reads were mixed with simulated human reads corresponding to approximately 1× human-genome coverage to model host-dominated, low-microbial-biomass samples. Reads were 150 bp with Illumina HiSeq 2500 sequencing profile. For the second benchmark, we used the 20 simulated samples from the CAMI III human-gut toy metagenome and evaluated using the corresponding CAMI gold-standard profiles obtained from the Critical Assessment of Metagenome Interpretation (CAMI) benchmarking portal (16). There are 20 samples containing 33.27 million reads on average. The CAMI III human-gut toy dataset also contains viral, fungal and plasmid sources, but this benchmark focused on profiling of bacterial genome abundance. Shotgun metagenomic sequencing data from the Canadian enteric pathogen cohort described by Peterson et al. (2022) were obtained from NCBI BioProject PRJNA786578 (18). It contains stool metagenomes with matched laboratory results for Salmonella and Campylobacter and was used for benchmarking detection of pathogens from metagenomes. Culture, PCR, and composite culture-or-PCR definitions were evaluated, with culture used as the primary reference standard. All 322 sequencing runs with matched laboratory results were analyzed as deposited (303 distinct specimens).

### Reference databases and software benchmarked

Two whole-genome reference databases were used to evaluate the effect of database choice. The GTDB database comprised bacterial and archaeal species-representative genomes from release R226. The RefSeq database was constructed from the NCBI RefSeq release 236, using the NCBI reference genome for each species (one genome per species). Winnow-tax, sylph, and Kraken2/Bracken were evaluated using both GTDB and RefSeq reference databases. KMCP was evaluated using its GTDB database, whereas MetaPhlAn 4 used its own marker-gene reference database that reports NCBI taxonomy. Predictions were scored against truth represented in the corresponding taxonomic nomenclature. For the CAMI III RefSeq comparison, predictions were paired with the gold standard on NCBI species taxid rather than on species name. GTDB predictions were not directly compared with NCBI-named truth. For CAMI III, the NCBI gold-standard profile includes taxa that cannot be represented by species-level RefSeq predictions (e.g., placeholder strain designations). All RefSeq results reported here are scored against the complete truth, so such taxa are counted as missed species for every RefSeq-based method (Figures 5 and 6). Detection was scored per sample as precision, recall and F1 against the truth species set for that sample. Per-sample F1 on CAMI III was compared between winnow-tax and each comparator using two-sided paired t-tests (n = 20). Abundance accuracy was scored as the L1 distance between the predicted and the true relative abundance vectors, each normalised to sum to one within a sample across the union of truth and predicted species. Abundance error was computed on read-share abundance for the decreasing-coverage series, with genome-copy outputs (winnow-tax, sylph, MetaPhlAn 4) converted using genome length, and on genome-copy abundance for CAMI III, with read-based outputs (Kraken2/Bracken) divided by genome length. Where a method reported no species in a sample, recall and F1 score were scored as zero, precision was treated as undefined, and the L1 distance was omitted from the reported means.

**Figure 6.**
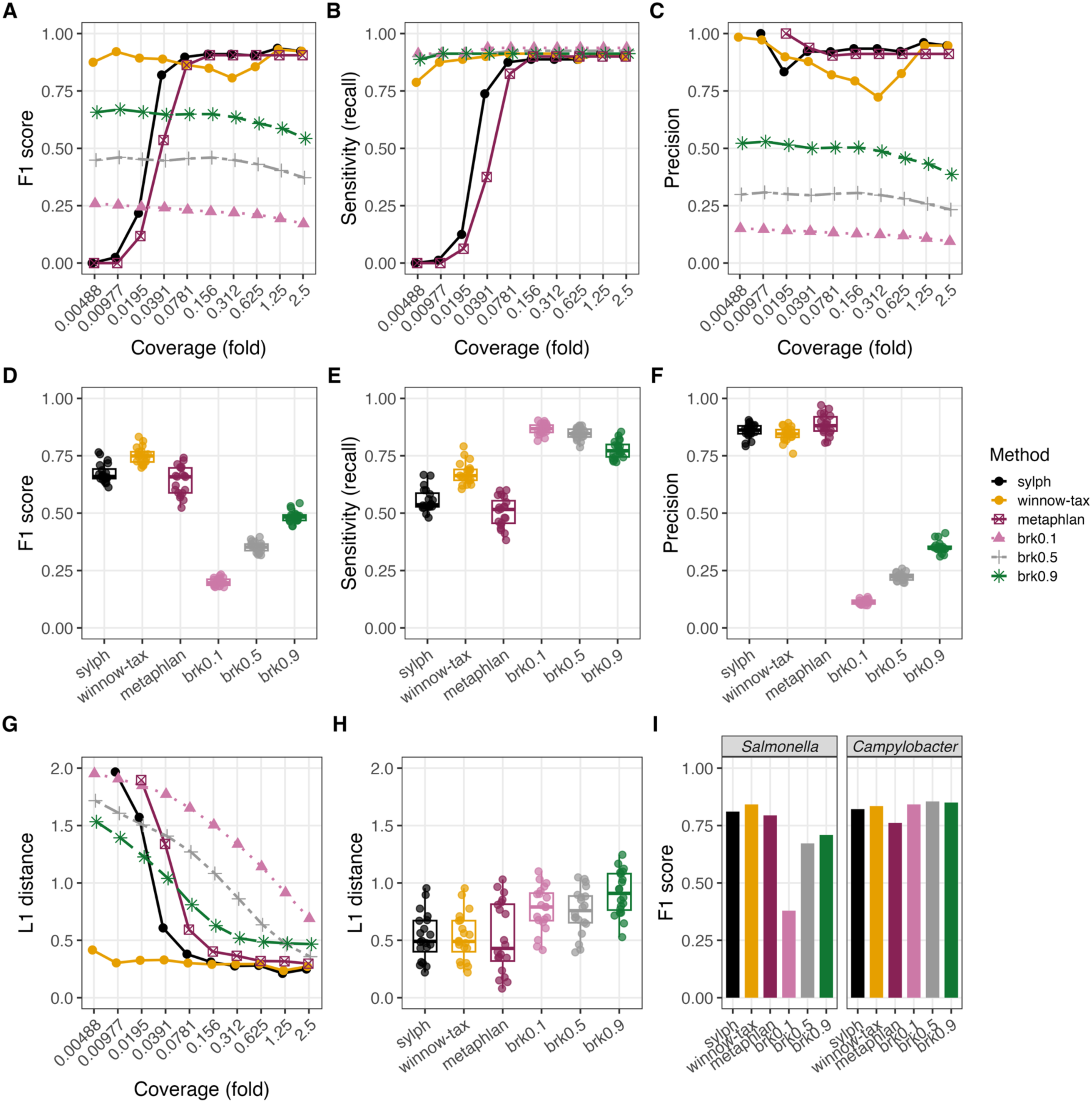
Comparison of winnow-tax with MetaPhlAn 4 in RefSeq/NCBI space. MetaPhlAn 4 was added to the RefSeq benchmark. (A-C) 512-fold coverage F1, sensitivity, and precision; (D-F) corresponding CAMI III metrics; (G-H) L1 distance; and (I) clinical pathogen F1. Winnow-tax achieved higher F1 primarily through increased recall, while MetaPhlAn 4 and sylph maintained similar precision but were more recall-limited at low coverage. MetaPhlAn 4 and winnow-tax achieved similar CAMI III L1 distances (0.524 and 0.528). Sylph reached 0.529 on the same measure, after its RefSeq profile was converted from sequence abundance to the genome-copy units of the gold standard (Methods). In the L1 panels (G and H), samples in which a method reported no species are omitted; in panel G this removes one sample for sylph and two for MetaPhlAn 4. brkX indicates Bracken applied to Kraken2 classifications at confidence X.

For the Kraken2 and Bracken benchmarks, reads were classified with Kraken2 (v2.17.1) against the GTDB R226 and RefSeq databases at confidence thresholds of 0.0, 0.1, 0.3, 0.5, 0.7, and 0.9. Results are shown for confidence 0.1, 0.5 and 0.9; confidence 0 produced near-zero precision. Species-level abundances were estimated from each confidence-specific report using Bracken v3.0.1. For Kraken2 and Bracken, a species was considered detected when at least 10 reads were assigned to it, consistent with Bracken’s default minimum-read threshold, to limit spurious low-count calls. MetaPhlAn (v4.2.2) was run using the vJan25 CHOCOPhlAn SGB marker database (mpa_vJan25_CHOCOPhlAnSGB_202503) with default parameters. Species-level profiles were used for downstream comparisons. Sylph (v0.9.0) was evaluated against both GTDB R226 and RefSeq. Reference sketches were generated at a subsampling rate of 200. Reads were sketched with sylph sketch and profiled with sylph profile using the default containment and ANI thresholds. KMCP v0.9.5 was evaluated against GTDB R226. The reference database was generated using kmcp compute with default parameters. Reads were searched using kmcp search. Taxonomic profiles were generated using all five predefined KMCP profiling modes and summarized at species level using GTDB taxonomy. Modes 1, 3, and 5 were used in the primary comparisons to represent permissive, intermediate, and more stringent filtering settings.

## Declarations

### Availability of data and material

Winnow-tax pipeline is available on GitHub (https://github.com/ssun6/winnow-tax). The CAMI III human-gut toy metagenome is available from the CAMI benchmarking portal. The clinical enteric metagenomes are available from NCBI BioProject PRJNA786578. Simulated reads for the decreasing-coverage series and the truth tables are available upon request. Additional requests and questions can be addressed to SS.

### Authors’ contributions

AAF and SS conceived the project. SS performed the analyses. AAF supervised the project. SS and AAF contributed to the writing of the manuscript. All authors read and approved the final manuscript.

## Reference

1. Kennedy KM, de Goffau MC, Perez-Munoz ME, Arrieta MC, Backhed F, Bork P, et al. Questioning the fetal microbiome illustrates pitfalls of low-biomass microbial studies. Nature. 2023;613(7945):639–49.

2. Guccione C, Patel L, Tomofuji Y, McDonald D, Gonzalez A, Sepich-Poore GD, et al. Incomplete human reference genomes can drive false sex biases and expose patient-identifying information in metagenomic data. Nat Commun. 2025;16(1):825.

3. Poore GD, Kopylova E, Zhu Q, Carpenter C, Fraraccio S, Wandro S, et al. Microbiome analyses of blood and tissues suggest cancer diagnostic approach. Nature. 2020;579(7800):567–74.

4. Tan CCS, Ko KKK, Chen H, Liu J, Loh M, Consortium SGKH, et al. No evidence for a common blood microbiome based on a population study of 9,770 healthy humans. Nat Microbiol. 2023;8(5):973–85.

5. Blanco-Miguez A, Beghini F, Cumbo F, McIver LJ, Thompson KN, Zolfo M, et al. Extending and improving metagenomic taxonomic profiling with uncharacterized species using MetaPhlAn 4. Nat Biotechnol. 2023;41(11):1633–44.

6. Shaw J, Yu YW. Rapid species-level metagenome profiling and containment estimation with sylph. Nat Biotechnol. 2025;43(8):1348–59.

7. Wood DE, Lu J, Langmead B. Improved metagenomic analysis with Kraken 2. Genome Biol. 2019;20(1):257.

8. Lu J, Breitwieser FP, Thielen P, Salzberg SL. Bracken: estimating species abundance in metagenomics data. PeerJ Comput Sci. 2017;3.

9. Nasko DJ, Koren S, Phillippy AM, Treangen TJ. RefSeq database growth influences the accuracy of k-mer-based lowest common ancestor species identification. Genome Biol. 2018;19(1):165.

10. Ye SH, Siddle KJ, Park DJ, Sabeti PC. Benchmarking Metagenomics Tools for Taxonomic Classification. Cell. 2019;178(4):779–94.

11. Breitwieser FP, Baker DN, Salzberg SL. KrakenUniq: confident and fast metagenomics classification using unique k-mer counts. Genome Biol. 2018;19(1):198.

12. Shen W, Xiang H, Huang T, Tang H, Peng M, Cai D, et al. KMCP: accurate metagenomic profiling of both prokaryotic and viral populations by pseudo-mapping. Bioinformatics. 2023;39(1).

13. Wright RJ, Fisher BR, Comeau AM, Langille MGI. From classification to confirmation: verifying taxonomic classifications by mapping metagenomic reads to reference genomes. Microb Genom. 2026;12(5).

14. Lander ES, Waterman MS. Genomic mapping by fingerprinting random clones: a mathematical analysis. Genomics. 1988;2(3):231–9.

15. Meslier V, Quinquis B, Da Silva K, Plaza Onate F, Pons N, Roume H, et al. Benchmarking second and third-generation sequencing platforms for microbial metagenomics. Sci Data. 2022;9(1):694.

16. Meyer F, Robertson G, Deng ZL, Koslicki D, Gurevich A, McHardy AC. CAMI Benchmarking Portal: online evaluation and ranking of metagenomic software. Nucleic Acids Res. 2025;53(W1):W102–W9.

17. Chaumeil PA, Mussig AJ, Hugenholtz P, Parks DH. GTDB-Tk v2: memory friendly classification with the genome taxonomy database. Bioinformatics. 2022;38(23):5315–6.

18. Peterson CL, Alexander D, Chen JC, Adam H, Walker M, Ali J, et al. Clinical Metagenomics Is Increasingly Accurate and Affordable to Detect Enteric Bacterial Pathogens in Stool. Microorganisms. 2022;10(2).

19. Meyer F, Fritz A, Deng ZL, Koslicki D, Lesker TR, Gurevich A, et al. Critical Assessment of Metagenome Interpretation: the second round of challenges. Nat Methods. 2022;19(4):429–40.

20. Sczyrba A, Hofmann P, Belmann P, Koslicki D, Janssen S, Droge J, et al. Critical Assessment of Metagenome Interpretation-a benchmark of metagenomics software. Nat Methods. 2017;14(11):1063–71.

21. McIntyre ABR, Ounit R, Afshinnekoo E, Prill RJ, Henaff E, Alexander N, et al. Comprehensive benchmarking and ensemble approaches for metagenomic classifiers. Genome Biol. 2017;18(1):182.

22. Wright RJ, Comeau AM, Langille MGI. From defaults to databases: parameter and database choice dramatically impact the performance of metagenomic taxonomic classification tools. Microb Genom. 2023;9(3).

23. Chiu CY, Miller SA. Clinical metagenomics. Nat Rev Genet. 2019;20(6):341–55.

24. Gu W, Miller S, Chiu CY. Clinical Metagenomic Next-Generation Sequencing for Pathogen Detection. Annu Rev Pathol. 2019;14:319–38.

25. Breitwieser FP, Lu J, Salzberg SL. A review of methods and databases for metagenomic classification and assembly. Brief Bioinform. 2019;20(4):1125–36.

26. Smith RH, Glendinning L, Walker AW, Watson M. Investigating the impact of database choice on the accuracy of metagenomic read classification for the rumen microbiome. Anim Microbiome. 2022;4(1):57.

27. Parks DH, Chuvochina M, Chaumeil PA, Rinke C, Mussig AJ, Hugenholtz P. A complete domain-to-species taxonomy for Bacteria and Archaea. Nat Biotechnol. 2020;38(9):1079–86.

28. Schoch CL, Ciufo S, Domrachev M, Hotton CL, Kannan S, Khovanskaya R, et al. NCBI Taxonomy: a comprehensive update on curation, resources and tools. Database (Oxford). 2020;2020.

29. Chen S, Zhou Y, Chen Y, Gu J. fastp: an ultra-fast all-in-one FASTQ preprocessor. Bioinformatics. 2018;34(17):i884–i90.

30. Li H. Minimap2: pairwise alignment for nucleotide sequences. Bioinformatics. 2018;34(18):3094–100.

31. Huang W, Li L, Myers JR, Marth GT. ART: a next-generation sequencing read simulator. Bioinformatics. 2012;28(4):593–4.

